# Evolution of local mutation rate and its determinants

**DOI:** 10.1101/054825

**Authors:** Nadezhda V. Terekhanova, Vladimir B. Seplyarskiy, Ruslan A. Soldatov, Georgii A. Bazykin

## Abstract

Mutation rate varies along the human genome, and part of this variation is explainable by measurable local properties of the DNA molecule. Moreover, mutation rates differ between orthologous genomic regions of different species, but the drivers of this change are unclear. Here, we compare the local mutation rates of several species. We show that these rates are very similar between human and apes, implying that their variation has a strong underlying cryptic component not explainable by the known genomic features. Mutation rates become progressively less similar in more distant species, and these changes are partially explainable by changes in the local genomic features of orthologous regions, most importantly, in the recombination rate. However, they are much more rapid, implying that the cryptic component underlying the mutation rate is more ephemeral than the known genomic features. These findings shed light on the determinants of mutation rate evolution.

---

Mutation rate is known to vary substantially between chromosomal regions^1–3^, and this variation is medically relevant^4–7^. Mutagenesis is affected by a range of biochemical processes, most importantly, meiotic recombination, replication and transcription, as well as by chromatin structure. Consistently, the local mutation rate (LMR) is strongly dependent on genomic features such as replication timing^8^, density of DNase-hypersensitive sites (DHSs)^9^, gene density, histone modifications, GC-content etc.^10–12^. Still, up to 70% of the human germline LMR variation at megabase scale cannot be explained by the known features^11^. Understanding the LMR variation and its causes is critical for inferring genomic functional elements and the genetic basis of heritable disease and cancer^4,7,13^.

The LMR landscape is dynamic. LMRs co-vary between closely related species^14^, but are almost independent of each other in remote species^15^. While the variation in LMR has been studied extensively, the dynamics and causes of LMR evolution are poorly understood (but see refs. ^2,12,14^. LMR evolution may be driven by changes in known genomic features or by other factors.

Evolution and co-evolution of the LMR and genomic features can be studied by analyzing the correlations between LMRs and features of orthologous genomic regions in species at a range of phylogenetic distances from each other. Here, we make use of complete genome alignments of 9 primate species, and of mouse, to study the evolution of the LMR between closely related vertebrates. We show that although only less than a half of the variance in LMR either in human or in apes can be explained by the known human genomic features, the LMRs in human and in apes are very strongly correlated, implying the existence of a strong “cryptic” component of the LMR variability. Furthermore, most of the genomic features are evolutionally stable and are good predictors of the LMR even in distantly related species; still, some changes in the LMR between species may be traced to changes in the underlying features, notably, in the recombination rate. By contrast, the “cryptic” fraction of the LMR variation not explainable by genomic features evolves very rapidly.

## RESULTS

### LMRs are strongly correlated between humans and apes

We study the multiple sequence alignment of 8 primate genomes (chimpanzee, gorilla, orangutan, gibbon, rhesus macaque, green monkey, squirrel monkey and marmoset) with human^16^, split into 2,261 1Mb non-overlapping windows (the results obtained for 100Kb windows were generally similar; Supplementary Note 1), together with the data on human polymorphism and de *novo* mutations in these same windows. To minimize the effect of selection on LMR estimates, we exclude exons and UTRs, and include the mean frequency of minor allele (MAF) in non-coding windows among the analyzed genomic features (see below). Still, selection acting at non-coding regions may confound inference of mutation rate variability (see Discussion).

For each species, we infer the nucleotide substitutions since its divergence from the last common ancestor with its closest relative (Fig. 1e,f), and count the number of such substitutions in each window as a proxy for the LMR (divergence-based LMR, dLMR). Additionally, for humans, we also estimate LMR using the numbers of rare SNPs^17^ (polymorphism-based LMR, pLMR) and the numbers of observed *de novo* mutations ^18^ (mLMR). While mLMR is the gold standard for LMR measurements, the data on it is limited (Supplementary Note 2), and we only use it for validation. pLMR is slightly better correlated with mLMR than dLMR, although both correlations are significant (P<0.01; Supplementary Fig. 1). We exclude substitutions prone to biased gene conversion from analyses (Methods and Supplementary Note 3).

The LMRs are strongly correlated between human and chimpanzee (R^2^=0.82 for dLMR, P < 2.2×10^−16^; R^2^=0.46 for pLMR, P < 2.2×10^−16^; Fig. 1a-b). When less related species are considered, this correlation decays with phylogenetic distance, reaching the minimal value among primates in marmoset (R^2^=0.27 for dLMR, P < 2.2×10^−16^; R^2^=0.04 for pLMR, P < 2.2×10^−16^; Fig. 1a-b), and is even lower in mouse (R^2^=0.11 for dLMR, P < 2.2×10^−16^; R^2^=0.02 for pLMR, P < 3.13×10^−7^; Supplementary Fig. 2). This decay is independent of the decrease in the fraction of alignable nucleotides with phylogenetic distance, as the correlation between LMRs remains similar when only the columns of the multiple alignments without gaps or ambiguous nucleotides in any of the species are considered (Supplementary Fig. 3). The proportion of the variance in LMR explainable by the human LMR decays by half at phylogenetic distance of ~0.04 substitutions per site, or ~16 million years^19^, roughly corresponding to the last common ancestor of human and orangutan. Human mLMR is also better correlated with the dLMRs of the more closely related species, compared with more distant ones (Supplementary Fig. 1 and 4).

**Fig. 1.**
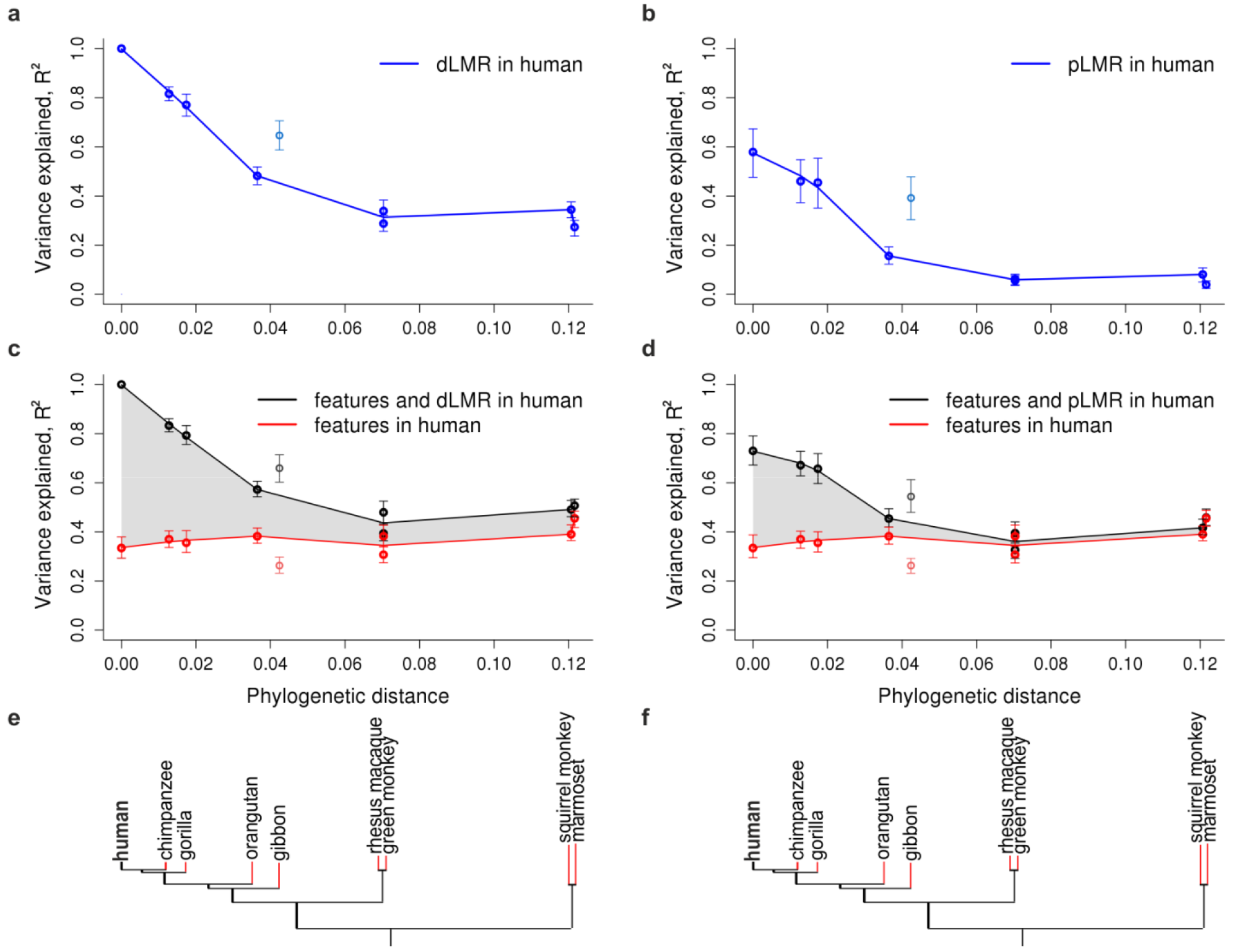
Local mutation rate (LMR) variation in primate species explained by genomic features and LMR in the human lineage. a-d.

For each primate species, the fraction of the explained variance in LMR (R^2^, vertical axis) is plotted against the phylogenetic distance from human (horizontal axis). The values for the gibbon, which has a high rate of rearrangements^21^, are plotted, but were not included in the fit. **a, b,** Variance in dLMR explained by the human dLMR (a) or pLMR (b). **c, d,** Variance in dLMR explained by human genomic features alone (red) or in combination with the human dLMR (c) or pLMR (d; black). The following features were included in the model: GC-content, recombination rate, number of exonic nucleotides, replication timing, number of DHSs, densities of H3K27ac, H3K27me3 and H3K9me3 histone marks and MAF. The shaded area represents the inferred fraction of the variance in non-human LMR explainable by the human LMR independently of the genomic features. Error bars correspond to 95% confidence intervals obtained by bootstrapping. **e, f,** Phylogenetic tree of the considered species. Red color denotes branches for which the LMR was calculated.

### A cryptic component to the LMR

The LMR depends on DNA properties^10–12^. The linear model that predicts the human dLMR from the measured genomic features of embryonic stem cells explains 33% of the variance in dLMR, which is in agreement with the previous estimates^11^ based on a feature annotation from a different tissue (Supplementary Note 4). The remaining variance may be random, or associated with genomic features not picked up by our analyses. The fact that the human LMR is a good predictor for the LMR in apes, in particular, in chimpanzee and in gorilla (Fig. 1a-b), suggests that the LMR variation not explainable by the measured genomic features still has a strong nonrandom component conserved between species.

To better understand this cryptic component, we first ask how well the human genomic features predict the LMR in non-human primates (Supplementary Fig. 5). For this, we construct, for each non-human species, a linear model predicting the dLMR in this species from human features alone and in combination with the human dLMR or pLMR. The non-human dLMR can be predicted nearly as well as the human dLMR by the features of the human orthologous segments (Fig. 1c-d, red line). This is consistent with the generally conservative nature of genomic features^20^. Indeed, most of the features are very strongly correlated even between human and mouse (Table 1).

**Table I.** Correlations between human and mouse genomic features in 1Mb genomic windows.

| **Feature** | **Pearson's R** | **P-value** |
| --- | --- | --- |
| LMR | 0.34 (0.29, 0.38) | $<2.2 \times 10^{-16}$ |
| Recombination rate | 0.01 (-0.04, 0.06) | 0.67 |
| DHSs | 0.83 (0.82, 0.85) | $<2.2 \times 10^{-16}$ |
| Replication timing | 0.71 (0.69, 0.74) | $<2.2 \times 10^{-16}$ |
| GC-content | 0.94 (0.93, 0.94) | $<2.2 \times 10^{-16}$ |
| Exonic nucleotide density | 0.95 (0.94, 0.95) | $<2.2 \times 10^{-16}$ |
| H3K9me3 | 0.32 (0.28, 0.37) | $<2.2 \times 10^{-16}$ |
| H3K27ac | 0.73 (0.70, 0.75) | $<2.2 \times 10^{-16}$ |
| H3K27me3 | 0.66 (0.63, 0.69) | $<2.2 \times 10^{-16}$ |

By contrast, adding the human LMR to the linear model radically increases the fraction of explained variance (Fig. 1c-d, black line): from R^2^=0.37 to 0.67 (for pLMR) or to 0.83 (for dLMR) in chimpanzee, and from 0.36 to 0.66 (for pLMR) or to 0.79 (for dLMR) in gorilla. Therefore, in closely related apes, the linear model that includes both the data on genomic features and the human polymorphism or divergence data explains about twice as much variance in LMR as the model with the genomic features alone, implying that about a half of the explained variance in LMR is cryptic (Fig.1c-d, shaded area).

Furthermore, there is a striking difference in how the explanatory power of genomic features and LMR changes with phylogenetic distance. While the human genomic features explain about as much variance in the LMR for distantly related as for closely related species, the human LMR predicts the LMR in closely related apes much better than that in less related primates. Therefore, unlike the measured genomic features which are relatively stable, the cryptic component of variation in LMR is short-lived. For the mouse LMR, human genomic features are much better predictors than the human LMR (Supplementary Fig. 2), suggesting that conserved features are important predictors of the LMR at large phylogenetic distances, while the LMR of a remote species carries no additional information. The cryptic component decays uniformly with phylogenetic distance, with the exception of the gibbon genome which carries an unusually high number of rearrangements^21^. Again, the shape of this decay is independent of the differences in alignment quality between species (Supplementary Fig. 3).

### Stability of genomic features in determination of the LMRs

The LMRs of orthologous genomic regions evolve with time (Fig. 1). The fraction of the variance in LMR explained by the human genomic features is similar in closely related and in distantly related primate species. Still, it is possible that individual features, and thus their power to predict the LMR in another species, change at different rates. We asked to what extent changes in LMR are determined by the evolution of individual features. As the data on genomic feature landscapes of primates is limited^22,23^, we addressed this question indirectly.

For this, we estimated the fraction of the variance in dLMR explained by individual human features. Because features are correlated with each other (Supplementary Fig. 6), we performed the ANOVA type III analysis to single out the independent contribution of each feature accounting for the contributions of other features (Fig. 2a). The estimated relative contributions of different features to the human dLMR are in line with previous work^10,11,14^. For the dLMRs in non-human primates, they are also mostly similar to those for the human dLMR, and for most features are independent of the phylogenetic distance to the analyzed species (Supplementary Fig. 7). The only feature with contribution declining with the phylogenetic distance is the recombination rate (P=0.009 for the correlation between R^2^ and the phylogenetic distance, Supplementary Fig. 7). The variance explained by it is high initially, in line with the evidence for its major effect on LMR^24–26^; but decreases rapidly with phylogenetic distance, from 6.02% for humans to 0.0005% for marmoset (Fig. 2a). This decay is linked to recombination *per se,* rather than to the associated process of gene conversion, as our analysis excludes the substitutions prone to biased gene conversion. Instead, they are in agreement with the gBGC-independent mutagenic role of recombination^26,27^. The observed decay suggests that changes in recombination are rapid, in line with its known high evolvability^28–30^. Such changes may contribute to changes in the LMR.

**Fig. 2.**
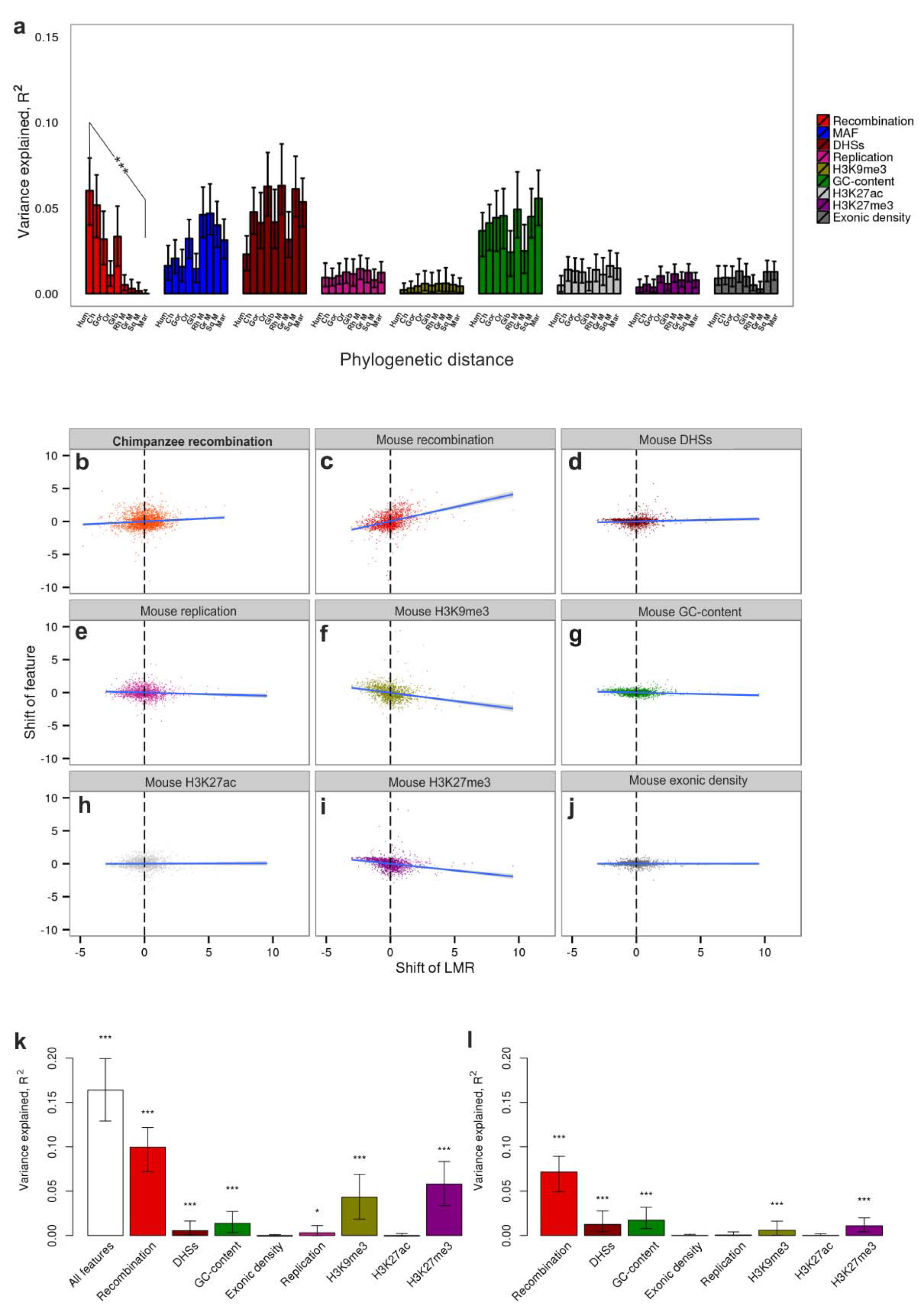
LMR explained by individual genomic features. a.

Variance of the LMR explained by the genomic features. For each genomic feature, the fraction of the variance explained by this feature in ANOVA type III analysis is shown for each primate at increasing phylogenetic distances from human. The order of the species along the horizontal axis is the same as in Figure 1 The asterisks indicate the significance of the negative correlation between the R^2^ and the phylogenetic distance. **b-j**, Scatterplots for raw correlations of the changes in the LMR and shifts of genomic features maps between human and mouse lineages. Each dot corresponds to a 1 Mb window. **k** and **l**, Changes in the LMR explained by the changes in genomic features between human and mouse. Vertical axis, fraction of variance in differences in LMRs explainable by differences in genomic features between human and mouse. Columns correspond to the variance explained by features measured by R^2^ (**k**) or ANOVA type III analysis (**l**). Error bars correspond to 95% confidence intervals obtained by bootstrapping. Asterisks indicate the significance of the deviation of the regression line from 0 (*: P<0.05, **: P<0.01, ***: P<0.001).

### Changes in recombination rate are associated with changes in LMRs

The importance of the local recombination rate for the LMR evolution is further supported by comparisons with chimpanzee and mouse. For these two species, recombination maps are available^30,31^. Recombination is poorly conserved between species: the human recombination rate is only weakly correlated even with that of chimpanzee (R^2^=0.24, P < 2.2×10^−16^), and not correlated with that of mouse (Table 1).

For each genomic window, we compared the interspecies differences in dLMR with differences in recombination rates. In both human-chimpanzee and human-mouse comparisons, they were weakly positively correlated (R^2^=0.01, P < 7×10^−6^ for chimpanzee, Fig. 2b; and R^2^=0.1, P<2.2× 10^−16^ for mouse; Fig. 2c), implying that an increase in the recombination rate of a genomic region between species is associated with an increase in LMR, and vice versa.

For the human-mouse comparison, we also analyzed several other genomic features, asking whether their changes are correlated with changes in the LMR. In total, ~16.4% of the variance in dLMR differences between branches could be explained by differences in the feature landscapes (Fig. 2k). When contributions from individual variables were considered, differences in recombination rate alone explained ~10% of the variance (Fig. 2c,k), while other features explained substantially less (Fig. 2d-j, k). To single out the genomic features in which changes between mouse and human lineages independently contribute to changes in the LMR between these two species, we performed the ANOVA (type III) analysis (Fig. 2l). Changes in only a few of the genomic features significantly contributed to changes in the dLMR. The most substantial contributor was recombination. Although the recombination landscape changes rapidly, and human recombination hotspots are not informative about the positions of hotspots in mouse (Table 1), changes in recombination rate landscape explain changes in dLMR more than those of any other features.

To better understand the link between changes in recombination and mutation, we studied the genomic windows in which the LMR has been substantially accelerated or decelerated in the human lineage, or in the chimpanzee lineage, since divergence from the human-chimpanzee common ancestor. In the human-accelerated regions (HARs), the human recombination rate is substantially higher than the genome average (one-sided Wilcoxon rank sum test P=2.9×10^−9^, Fig. 3a), implying that the HARs frequently carry recombination hotspots. By contrast, in the chimpanzee-accelerated regions (CARs), the human recombination rate is only slightly higher than the genome average (P=8.0×10^−3^, Fig. 3b). Together, these data imply that the HARs are frequently associated with recombination hotspots that are short-lived, so that they increase the mutation rate in human more than in chimpanzee. Reciprocally, the chimpanzee recombination rate is elevated in CARs (P=3.7×10^−7^, Fig. 3d), slightly more than in HARs (P=7.4×10^−6^, Fig. 3c).

**Fig. 3.**
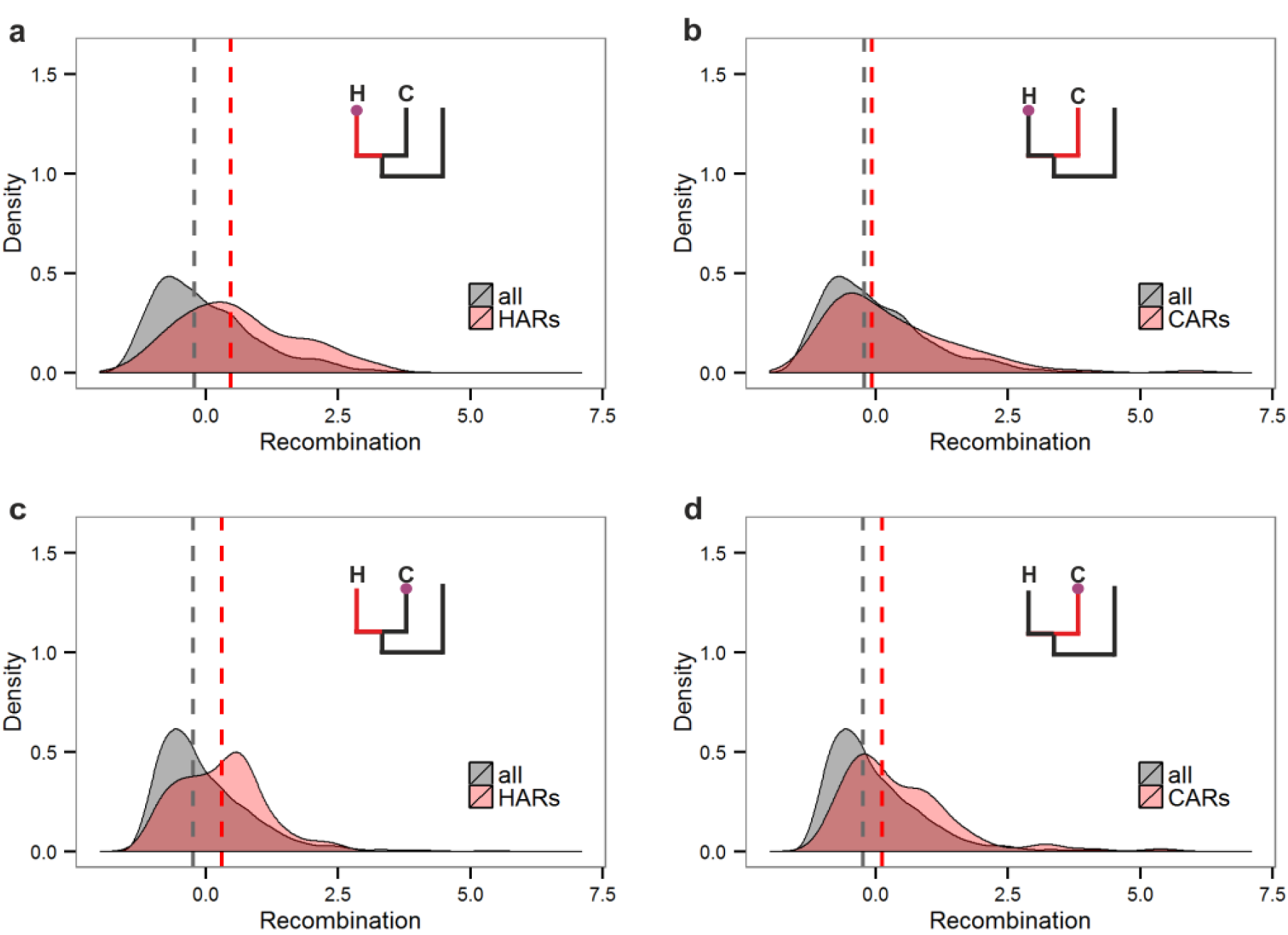
Distributions of human recombination scores (a-b) in HARs (a) and CARs(b), and of chimpanzee recombination scores (c-d) in HARs (c) and CARs (d).

The schematic phylogenies show the lineage in which the LMR was increased in red (H, human, or C, chimpanzee), and the species in which recombination was measured, as a circle. The dashed line corresponds to the median recombination rate in the ARs and all genomic regions.

We also analyzed the genomic regions that were substantially decelerated in human (human decelerated regions, HDR) or chimpanzee (CDR). The human recombination rate at HDRs as well as the chimpanzee recombination rate at CDRs were slightly reduced compared to the genome average (one-sided Wilcoxon rank sum test P= 7.7×10^−4^ and P= 3.9×10^−4^ respectively; Supplementary Fig. 8). This implies that the LMR deceleration is also partially caused by recombination-related factors, although likely to a smaller extent than LMR acceleration. All these patterns were associated with the recombination *per se* rather than biased gene conversion (Supplementary Note 5).

## DISCUSSION

While the LMR is known to differ between species ^2,32,33^, the rate and the driving forces of the LMR evolution are obscure. To our knowledge, this study is the first quantitative analysis to this end.

Our approach to estimation of the germline mutation rate from divergence and polymorphism data has three important caveats. First, selection and GC-biased gene conversion (gBGC) can affect both divergence and polymorphism. To limit the effect of selection, we only analyzed intronic and intergenic regions, as only ~8% of mutations at these regions are affected by selection^34^. The contribution of MAF to the explained variance is significant (Fig. 2a), implying that the effect of selection on LMR is still high even within the noncoding regions. However, the contribution of MAF is much smaller than the unexplained component of the LMR variance, implying that the high correlation between LMRs of closely related species is not due to common selection pressures. Moreover, the contribution of MAF is roughly constant between species at different phylogenetic distances (Fig. 2a), implying that selection pressures are rather stable, and their changes contribute little to the LMR evolution. gBGC is indeed associated with divergence^29^ and polymorphism; however, its effect is weak or absent in the considered subset of substitutions (Supplementary Notes 3 and 5). Second, the estimates of the LMR can be affected by the polymorphism in the ancestral population. In particular, the differences in coalescence times between genomic regions may inflate the correlation between the dLMRs estimated from two descendant sister branches^35–37^. Similarly, such differences may contribute to the association between the dLMR and the recombination rate, as genomic regions with higher recombination rates have larger local effective population size, and therefore longer coalescence times^37–39^. Such phenomena do not affect correlations with *de novo* mutations (Supplementary Fig. 1). They are also unlikely to contribute to pLMRs, as transspecies polymorphisms are rare^40,41^, and can only contribute to the correlations between dLMRs, and the correlation between LMR and recombination, of human, chimpanzee and gorilla - the species in which the contribution of the ancestral polymorphism to divergence is reported^42^. In all other comparisons, a relatively long period of independent divergence (black edges in the phylogeny of Fig. 1) separates the branches at which the dLMR is measured (red edges in Fig. 1). Furthermore, the time to coalescence of a genomic region is highly correlated with its exonic density^38^, and the correlation between the exonic density and the dLMR is low (R^2^=0.01, Fig. 2a), suggesting that the ancestral polymorphism also does not contribute much to correlations between dLMRs, or to correlations between the LMR and the recombination, even in the most closely related species. Still, such confounders may affect dLMR estimates to some degree; in particular, they may be the reason why the correlations between dLMRs are higher and more long-lived than the correlations between the pLMR in human and dLMRs in other species. Third, the mutation rate estimation is dependent on the window size, with small windows giving unreliable estimates^15^. The window sizes we use, 100Kb and 1 Mb, are a compromise between resolution and robustness.

Given these caveats, we show that the correlation between the LMRs of closely related species is surprisingly high. As a result, the LMR of a species can be much better predicted by the LMR of a closely related species than by its own genomic features. Still, the LMR is plastic, being rather poorly conserved even between moderately related species of primates, and evolves much faster than the known genomic features. These results imply the existence of a strong but transient cryptic component in LMR variation; the unknown genomic features that underlie it must undergo a rapid turnover, changing at the timescale of a few tens of millions of years.

By contrast, the predictive power of human genomic features for the non-human LMR changes little with time since divergence from human, probably due to the conservative nature of these DNA properties. As a result, in the human-mouse comparison, human genetic features are better predictors of the LMR in mice than the human LMR.

Our findings therefore imply that changes in the known genomic features may not be responsible for most of the changes in LMR in the course of evolution, at least at short timescales. Nevertheless, we can still measure the correlation between the genomic features and the LMR defined in another species, and trace how this correlation changes with distance between species, as a proxy for the rate of evolution of DNA landscape. A decrease in correlation between the LMR of one species and DNA properties of another species mirrors changes in genomic features with time.

Using this approach, we show that features differ in their contributions to the mutation rate dynamics. The recombination rate is known to be one of the key features that influence the mutation rate variation in humans^28,43,44,45^. Furthermore, local recombination rate is very plastic, and its hotspots vary dramatically even between closely related species^46,47^; the correlation of the recombination rates is low even between human and chimpanzee. Our results show that changes in the recombination rate are among the biggest contributors to the LMR evolution among the studied genomic features.

The high similarity of LMRs between closely related species implies that the estimates of the neutral mutation rate at a genomic region could be substantially improved by considering the mutation rate at homologous regions from closely related species. By contrast, the functional annotation may be more informative about the mutation rate in a more distantly related species.

More generally, existing approaches to inference of functional genomic elements often use interspecies conservation as a proxy for function. This involves two assumptions about the mutation process: first, that the mutation rate is uniform along the genome; second, that it is constant between species. Violations of these assumptions can lead to false inferences. The first assumption is now being relaxed^2,4,48^; however, the second largely remains in place. Appreciation of the importance of the variation in the genomic mutation rates in the course of evolution has revolutionized the field of molecular dating^2,49^. Analogously, we propose that understanding the LMR variation between species and its causes may help predict the likelihood of mutations and infer their functional importance.

## Online Methods

### Alignments

The multiple sequence alignment of 8 primate genomes (chimpanzee, gorilla, orangutan, gibbon, rhesus, green monkey, marmoset and squirrel monkey) with the hg19 version of the human genome assembly was obtained from the multiple alignment of 100 vertebrate species downloaded from the UCSC Genome Browser (genome.ucsc.edu/)^16^ using multiz-tba^50^. The alignment was then split into non-overlapping 1Mb or 100Kb windows. Exonic nucleotides, UTRs, repeats, ambiguous nucleotides and CpG dinucleotides were masked, and windows with more than 20% gaps and masked nucleotides in any of the nine species were excluded from further analysis. This procedure resulted in 2,261 1Mb windows, or in 23,551 100Kb windows. Independently, multiple alignments of the same 9 genomes with the mouse genome were obtained in the same way. Since there are more gaps in the primates-mouse alignment than in the primates-only alignment we excluded windows with more than 10% gaps and masked nucleotides in any of the species. This procedure resulted in fewer windows: 1,454 1Mb windows, and 16,449 100Kb windows.

To make sure that our results are not affected by differences in the fraction of excluded (unaligned or masked) nucleotides between genomic regions, we repeated the analyses using only those alignment columns where all species had an aligned and unmasked nucleotide, and only those windows where there were >10% in the primates-mouse alignment. This resulted in 1,091 1Mb windows for the primates-and-mouse alignment.

### Genomic features mapping

Replication time, DHSs and histone modifications H3K9me3, H3K27ac, H3K27me3 as measured by the ENCODE project^51^ were downloaded from the UCSC Genome Browser (genome.ucsc.edu/ENCODE/). The analysis in the main text uses these maps obtained for embryonic stem cells. Additionally, we used the maps for 5 other tissues: GM12878, HUVEC, NHEK, Hela-S3, K562 and (Supplementary Fig. 9, Supplementary Table 1). Recombination rates were obtained from the HapMap project (hapmap.ncbi.nlm.nih.gov/). gBGC tracts obtained using phastBias with the parameter B=3 were taken from ref. ^52^. The value of a feature for a genomic window was calculated as the weighted average of this feature, excluding masked nucleotides and gaps. MAFs and polymorphism data were obtained for all human SNPs except those that were W↔S from the 1000 genomes project^17^. Mean values of MAFs for each window were calculated. pLMRs were calculated as the polymorphism’ frequencies by utilizing only 50% of the rarest SNPs. The list of *de novo* mutations was obtained from the ref.^18^.

Chimpanzee and mouse recombination rates were obtained from refs. ^30^ and ^31^ respectively. Mouse DHSs, replication timing, and histone modifications H3K9me3, H3K27ac and H3K27me3 measured by the Mouse ENCODE project for ES cells (mouse genome version mm9) were downloaded from the UCSC Genome Browser (genome.ucsc.edu/ENCODE/), and mouse genomic coordinates were converted into hg19 coordinates using liftOver^16^.

The fraction of exonic nucleotides were calculated for all nucleotides (including masked ones) at each genomic window. To compare changes between features in the course of evolution, the values of each feature for each species were normalized to mean=0 and variance=1.

### Inference of LMR

As a proxy for the local mutation rate (LMR) in a genomic window, we inferred the number of single-nucleotide substitutions that occurred in this window at a given phylogenetic branch (red in Fig. 1e-f). For this, we inferred the ancestral states using maximum parsimony, and measured the fraction of substituted nucleotides among all unmasked non-gapped nucleotides. These fractions for all genomic windows were defined as LMRs normalized to mean=0 and variance=1, and LMR was defined as this normalized value. To avoid the confounding effect of GC-biased gene conversion (gBGC), we excluded W↔S substitutions (Supplementary Note 3). The analyses including the W↔S substitutions are presented in Supplementary Fig. 10-13. As an alternative approach, we also utilized the maximum likelihood as implemented in the baseml program of the PAML package^53^, using the REV model with no molecular clock; here, all substitutions, including W↔S, were analyzed, and the results were similar (Supplementary Figure 10).

We assessed the accuracy and power of LMR inference by bootstrapping nucleotide sites within each window. The observed dLMRs and pLMRs were very strongly correlated with the bootstrapped samples (dLMR: R=0.97, P< 2.2×10^−16^, pLMR: R=0.98, P< 2.2×10^−16^), implying that these estimates are robust. The correlation was weaker for mLMRs (R=0.75, P< 2.2×10^−16^; Supplementary Figure 14).

Since mutation rates differ between nucleotides^54^, our LMR estimates as described above may be confounded by the differences in the nucleotide composition of genomic windows. To address this, we additionally used an alternative procedure for LMR estimation that accounts for the nucleotide composition. We calculated, for each species *b* and genomic window *h,* the expected number of mutations Mbased on its nucleotide composition:

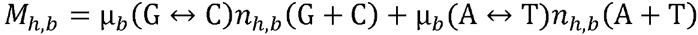

where μ_b_ is the genomic rate of the corresponding mutation in species *b;* and *n_h,b_* is the number of corresponding nucleotides in this window in species *b*. We then defined LMR as the ratio of the observed and expected numbers of mutations. This procedure yielded very similar results to those in the main text (Supplementary Figures 15-16).

### Explained variance of the LMR

Using linear regression as implemented in the lm function in R, we calculated the fraction of the variance in LMR between genomic windows in a species that can be explained by genomic features in the same and/or different species, and/or by the human LMR. Adjusted R^2^ values were used to minimize the effect of the number of explanatory variables. To calculate the contribution of each feature independently of the contributions of other features, we performed ANOVA type III tests using drop1 function in R. 95% CIs for R^2^ values were obtained by bootstrapping genomic windows in 200 bootstrap trials. Local polynomial regression fitting in Figure 1 was performed using the loess function of R.

### Correlating differences between LMR and genomic features

For each window, we calculated the difference in normalized LMR values between human and mouse or chimpanzee, and the differences in normalized feature values between the same two species. We then calculated Pearson's correlation coefficients between these values. We also performed ANOVA type III tests to estimate the independent contributions of changes in different genomic features to changes in the mutation rate, explaining changes in LMR between species with the drop1 function corresponding to changes in each feature separately. 95% CIs for R values were obtained by bootstrapping genomic windows in 200 bootstrap trials.

### Correlations with phylogenetic distance

To estimate the significance of the correlation between the phylogenetic distance and the R^2^ values, we obtained the distribution of Spearman correlation coefficients by bootstrapping genomic windows in 10,000 bootstrapping trials.

### Identification of HARs, HDRs, CARs and CDRs

We aimed to select the genomic windows such that the human (chimpanzee) LMR was substantially increased or decreased, compared with the chimpanzee (respectively, human) LMR. In this section, raw LMR values were used, i.e., normalization to mean=0 and variance=1 was not applied. First, we predicted the expected LMR *u_b,h_(exp)* in phylogenetic branch *b* for each genomic window h, accounting for the mean LMR of this window across all branches 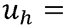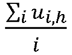 and the length of the branch leading to this species 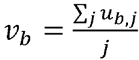 and normalizing by the mean LMR across all windows in all species:

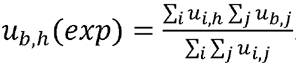

We compared this value for human or chimpanzee with the corresponding observed value of LMR, *u_b,h_(obs)*. To single out the genomic windows with LMR changes in the species of interest, we then selected all windows where the magnitude of change in this species sp_1_ (human or chimpanzee) was greater than the magnitude of change in the sister species sp_2_ (respectively, chimpanzee or human):

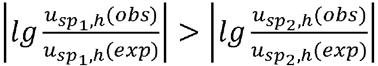

We ranked these windows by the magnitude of change in LMR in *sp_1_* compared with *sp_2_,*

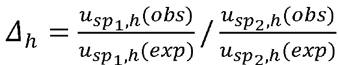
and identified accelerated regions (ARs) and decelerated regions (DRs) as the 100 windows with the highest and lowest values of *Δ_h_.* Summary statistics describing the properties of ARs and DRs are presented in Supplementary Table 2.

For all density plots, we used the kernel density estimation implemented in ggplots R-package with the default optimized bandwidth of smoothing (bw.nrd0). The scores for recombination and gBGC for all genomic windows were normalized to mean=0 and variance=1.

The 95% confidence intervals (asymptotic confidence intervals estimated based on Fisher's Z transform) are in parentheses.

## ACKNOWLEDGMENTS

We thank the members of Shamil Sunyaev's, Alexey Kondrashov's and Georgii Bazykin's labs for discussion and helpful comments on the manuscript. This work was supported by the Russian Foundation for Basic Research grant #15–34–21135–mol_a_ved to G.A.B. and by the Molecular and Cellular Biology Program of the Russian Academy of Sciences.

## AUTHOR CONTRIBUTIONS

NVT, VBS, RAS and GAB planned and designed the experiments, VBS and GAB directed the research, NVT analyzed the data, and RAS assisted in statistics and performed the power test analysis. All authors contributed to the preparation of the manuscript.

